# Enhancing slow-wave sleep via non-invasive brain stimulation modulates brain-to-blood clearance of Alzheimer’s disease biomarkers

**DOI:** 10.64898/2026.06.22.733841

**Authors:** Julia Ladenbauer, Paula Schümann, Youannes Rizk, Robert Malinowski, Buse Dikici, Antje Vogelgesang, Agnes Flöel

**Author notes:** **Correspondence:** Julia Ladenbauer; Agnes Flöel, Universitätsmedizin Greifswald, Klinik und Poliklinik für Neurologie, Ferdinand-Sauerbruch-Straße, 17475 Greifswald, Germany.

## Abstract

Sleep disturbances and neurodegeneration form a bidirectional vicious cycle. During slow-wave sleep, glymphatic processes are thought to facilitate the clearance of metabolic waste, including proteins implicated in Alzheimer’s disease (AD). Aging, and more prominently neurodegeneration, is associated with reductions in slow-wave activity (SWA), which may impair these processes. Within SWA, slow oscillations (SO, <1 Hz) and their coupling to sleep spindles capture key aspects of sleep microstructure. Non-invasive brain stimulation during sleep has emerged as a potential approach to modulate these dynamics; however, human evidence linking such modulation to AD biomarkers remains scarce.

In this exploratory mechanistic study, ten healthy older adults underwent one night of slow-oscillatory transcranial current stimulation (so-tDCS) and one sham night in a randomized crossover design, followed by five consecutive stimulation nights. Stimulation was applied during NREM sleep (N2/N3) in the first half of the night. Plasma phosphorylated tau (p-tau)181, β-amyloid (Aβ)42, Aβ40, and total tau were assessed overnight and longitudinally. EEG analyses quantified SO power and SO-spindle coupling. Due to the exploratory character of the study, analyses emphasized effect size estimation and explained variance.

So-tDCS induced small-to-moderate increases in SO power and SO-spindle coupling. Overnight increases in plasma p-tau181 were observed following stimulation relative to sham. Increases in SO power were strongly and positively associated with p-tau181 changes, explaining a substantial proportion of inter-individual variance. In contrast, shifts in SO-spindle coupling phase toward the SO up-state were associated with overnight increases in Aβ42 and Aβ40 and with longitudinal decreases in Aβ42 and the Aβ42/40 ratio.

Enhancing slow oscillatory dynamics during sleep is associated with changes in peripheral AD biomarkers. Differential associations for SO power and SO-spindle coupling timing suggest partially distinct links to tau and Aβ dynamics. These findings support sleep microstructure as a potential intervention target, and should be confirmed in larger cohorts.

## Background

Alzheimer’s disease (AD) is characterized by a decades-long preclinical phase during which pathological changes accumulate, rendering the identification of early risk factors and novel preventive strategies essential. Sleep disturbances frequently emerge years before the onset of cognitive symptoms and correlate with the accumulation of β-amyloid (Aβ) and tau proteins (1). The relationship between sleep and AD pathology is bidirectional: disrupted or fragmented slow-wave sleep (SWS) accelerates Aβ and tau accumulation (2–4), while Aβ and tau burden reciprocally reduce slow-wave activity (SWA) and SWS (5,6). Beyond these changes, AD pathology also disrupts other key oscillatory features of non-rapid eye movement (NREM) sleep, including a loss of sleep spindles and deterioration of the precisely timed slow oscillation-spindle coupling, which is widely implicated in memory consolidation (7–9). Critically, SWA and SWS decline progressively from midlife onward even in the absence of pathology (10,11), and cognitively healthy older adults frequently exhibit Aβ and tau deposits associated with elevated dementia risk (2).

One central mechanism through which sleep disruption may drive AD pathology involves the glymphatic clearance of neurotoxic proteins: accumulating evidence indicates that SWA facilitates the removal of Aβ and tau species from the brain (12). This process is thought to be mediated by the glymphatic system, a network of perivascular channels, glial cells, and cerebrospinal fluid (CSF) flux, which transports interstitial solutes out of the brain during deeper NREM sleep phases (13,14). Consistent with these findings, human studies demonstrated that a single night of uninterrupted sleep reduces Aβ and tau concentrations in CSF relative to sleep deprivation (15,16), whereas experimental disruption of SWA elevates overnight CSF Aβ concentrations (4).

Mechanistically, cortical slow-wave activity (SWA) during NREM sleep, particularly its high-amplitude, low-frequency component in the form of slow oscillations (SOs, < 1 Hz), is thought to drive this clearance by entraining rhythmic fluctuations in cerebral blood volume that promote CSF influx and interstitial solute efflux (17,18). This efflux is ultimately reflected in peripheral blood, as proteins cleared from the brain parenchyma into CSF are subsequently transported to peripheral blood, rendering plasma a scalable and minimally invasive index of brain clearance dynamics (16). Further supporting the relevance of sleep microstructure beyond SO activity, a pooled baseline analysis across three clinical trials found that precise SO-spindle coupling timing was a stronger predictor of plasma Aβ levels than SWA across healthy and cognitively impaired older adults (19), and improvements in SO-spindle coupling via acoustic stimulation were associated with later favorable changes in the plasma Aβ42/40 ratio in cognitively impaired older adults (20).

Despite this evidence, the precise contribution of the glymphatic system in humans remains a matter of ongoing debate (21,22); yet, there is broad consensus that SWS and its associated oscillatory activity are fundamental to the regulation of cerebral protein clearance during aging.

Building on this rationale, interventions aimed at enhancing SWA during SWS have been discussed as a promising preventive approach in healthy older adults and as a potential therapeutic strategy in those with cognitive impairment (23). One candidate intervention is slow-oscillatory transcranial current stimulation (so-tDCS), in which a weak oscillating electrical current (< 1 Hz) is applied to the scalp during NREM sleep. Previous work has shown that so-tDCS can augment endogenous slow oscillatory activity and improve the timed coordination between SO events and spindle activity in healthy older adults (24–26) as well as in individuals with MCI (27). Whether this approach can also influence brain clearance processes during sleep remains to be established.

We thus conducted an exploratory mechanistic study to examine whether enhancing SO activity via so-tDCS during sleep promotes the clearance of AD-relevant proteins, as reflected in peripheral blood concentrations. This study was designed to probe biological mechanisms and estimate effect sizes. We hypothesized that stimulation-induced increases in (<1 Hz) SWA would predict greater overnight protein clearance, evidenced by higher plasma Aβ42 and higher p-tau181 concentrations in the morning following so-tDCS compared to sham, and that repeated enhancement of SWA across five consecutive stimulation nights would be reflected in longitudinal shifts in plasma Aβ42, the Aβ42/40 ratio, and p-tau181 from before to after the intervention. Circadian and peripheral clearance confounds were controlled using fixed sampling times and fasting protocols. Additionally, baseline liver and kidney functions were assessed.

## Methods

### Participants

Eleven healthy older adults (aged 57- 82 years; mean 66.8 ± 6.7 years) were recruited at the Department of Neurology, University Medicine Greifswald, Germany (see Table 1 for baseline characteristics). Exclusion criteria included severe untreated neurological, psychiatric, or medical conditions; use of centrally active medication; manifest sleep disorders; alcohol or substance abuse; non-fluency in German; and MRI-detected brain pathologies (see Supplementary Methods for full list). One participant was excluded due to insufficient blood withdrawal at baseline, leaving ten participants for final analysis. Habitual sleep was monitored by actigraphy for five days prior to laboratory sessions; actigraphy and memory performance data are beyond the scope of the present paper and will be reported in a subsequent analysis.

**Table 1.**
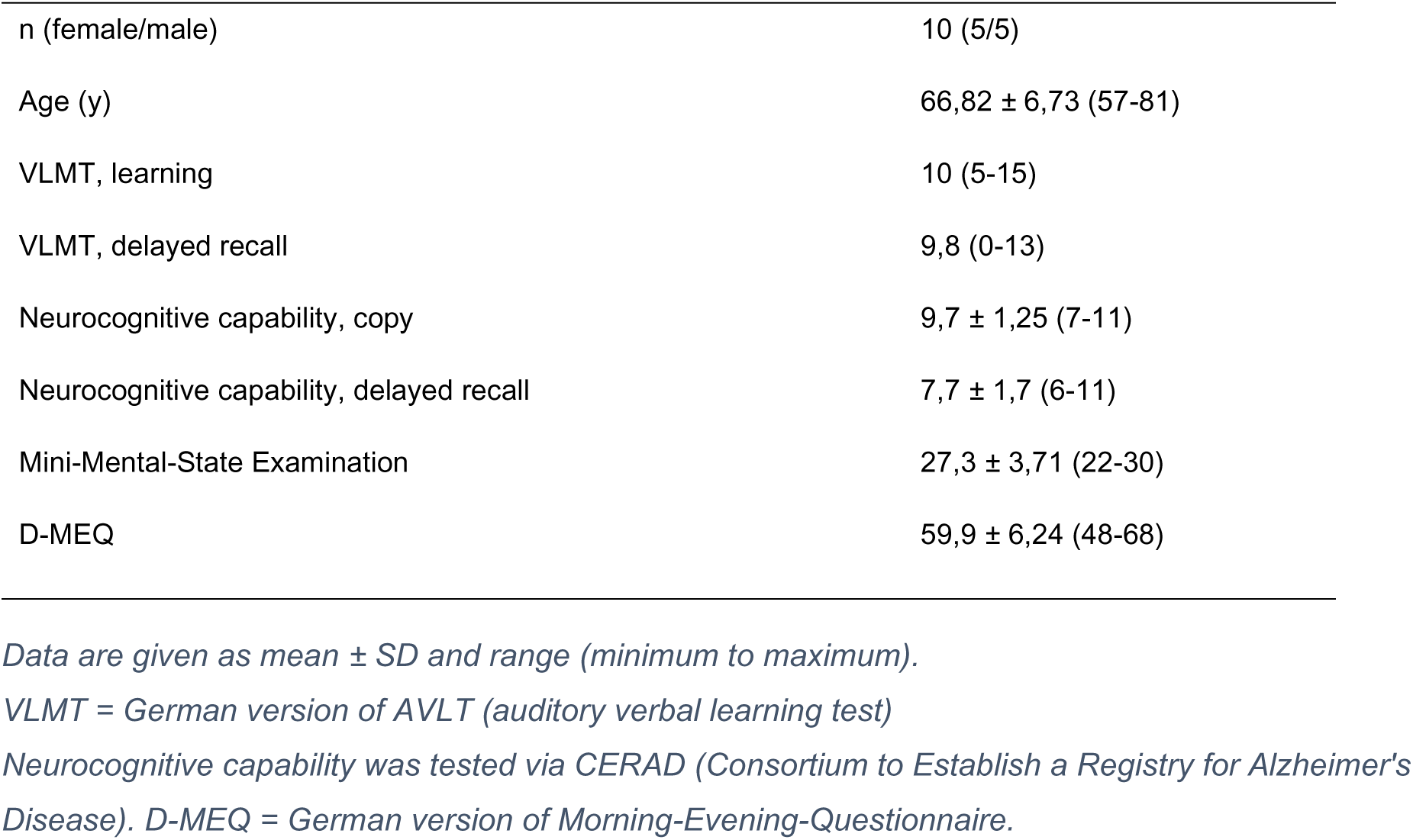
Baseline characteristics.

All participants provided written informed consent in accordance with the ethics committee of the University Medicine Greifswald and the Declaration of Helsinki and received an expense allowance.

### Study design and procedure

This pilot study investigated short- and longer-term effects of slow oscillatory transcranial current stimulation (so-tDCS) during NREM sleep on plasma AD biomarkers (p-tau181, Aβ42, Aβ40, total-tau) as surrogate markers of glymphatic clearance. As the temporal dynamics of potential stimulation effects on these recently validated plasma biomarkers could not be predicted a priori, two complementary analyses were pre-specified: (i) short-term effects: overnight biomarker changes under so-tDCS versus sham in a randomized counterbalanced crossover design (blood sampling at 22:00 and 08:00); and (ii) longer-term effects: morning plasma levels before versus after six consecutive stimulation nights (08:00, S1 vs. S9). To control for confounding influences, blood sampling times and fasting requirements (minimum 5 h) were kept constant across all sessions, and hepatic and renal function were assessed at baseline, as individual peripheral clearance capacity influences plasma biomarker levels (28).

All sessions were conducted at the sleep laboratory of the Department of Neurology, University Medicine Greifswald. Participants completed nine sessions across 13 days (Fig. 1). Session 1 (S1) comprised baseline neuropsychological assessment, blood sampling (08:00), memory testing (object-location and word pair task) and structural and resting-state MRI at a 3 T Siemens MAGNETOM Skyra scanner (Siemens Healthcare, Erlangen, Germany) to carefully characterize participants at baseline and to exclude pathologies. Habitual sleep was monitored by actigraphy for five days prior to laboratory sessions. S2 served as a laboratory adaptation night with full polysomnographic EEG recording. During S3 and S4, so-tDCS and sham stimulation were administered in a randomized counterbalanced crossover design; blood samples were collected before sleep (22:00) and upon waking (08:00). Five consecutive so-tDCS nights (S5- S9) followed to examine longer-term effects, with a final blood draw on the morning after S9 (08:00). Sampling times and fasting requirements (minimum 5 h) were kept constant across all sessions to control for circadian variation and individual differences in peripheral clearance. Actigraphy and memory performance data are beyond the scope of the present paper and will be reported in a subsequent analysis.

**Figure 1.**
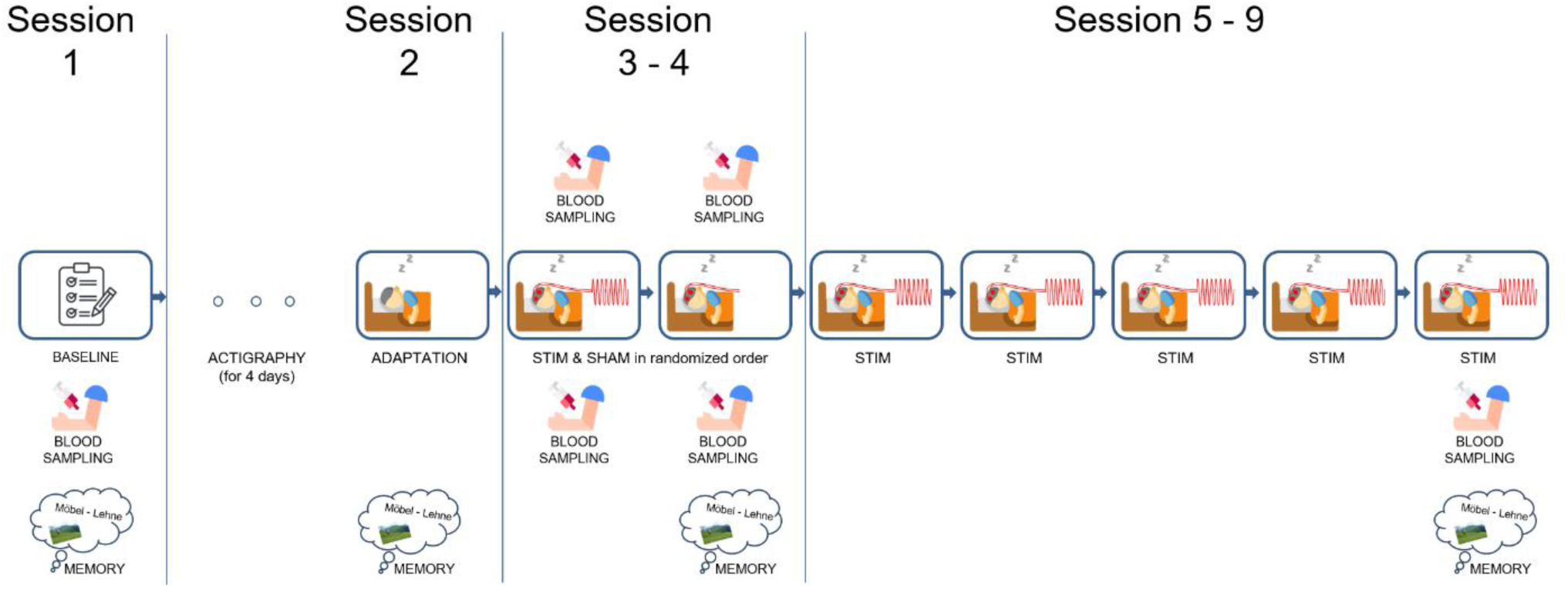
Study design overview. The study comprised nine sessions across 13 days. Session 1 included baseline neuropsychological assessment, blood sampling (08:00), memory testing and structural/resting-state MRI; participants subsequently wore an actigraphy device for five days to characterize habitual sleep. Session 2 served as a laboratory adaptation night with EEG recording and blood sampling (08:00) the following morning. Sessions 3- 4 constituted the crossover phase: participants received either slow oscillatory transcranial current stimulation (so-tDCS; 0.75 Hz anodal, repeated 5 min blocks during N2/N3 of the first half of the night) or sham stimulation in randomized counterbalanced order, with blood samples collected before sleep (22:00) and upon waking (08:00). Sessions 5- 9 comprised five consecutive so-tDCS nights to assess longer-term effects, with a final blood sample obtained on the morning after Session 9 (08:00). Memory outcomes will be reported separately.

### Brain stimulation

Stimulation electrodes (8 mm diameter) were placed bilaterally at frontal sites (F3, F4) with return electrodes on ipsilateral mastoids (M1, M2). Anodal current was applied via two battery-driven stimulators (DC-Stimulator, NeuroConn, Germany) modulated sinusoidally at 0.75 Hz (range 5-255 µA; DC baseline 130 µA; maximum current density 0.507 mA/cm²). Stimulation was initiated four minutes after stable N2 entry and delivered in 5-min blocks exclusively during N2/N3 sleep in the first half of the night (∼4 h), separated by stimulation-free intervals of at least 1.5 min; it was suspended if participants left N2/N3 and resumed after stable return for ≥1 min. During the sham condition, stimulation electrodes were placed identically to the so-tDCS condition, but the tCS device remained off. The sham session followed the same temporal pattern and procedural criteria as the so-tDCS condition.

### Plasma biomarker assessment

Blood (4 ml) was collected via brachial venipuncture at seven timepoints: five morning samples (08:00; S1- S4, S9) and two evening samples (22:00; S3, S4). Plasma was prepared according to SABB guidelines using K2EDTA tubes, centrifuged within 30 min (1800 × g, 10 min, 21°C), and immediately stored at −80° C until analysis. P-tau181 was quantified using the commercially available p-tau181 Advantage V2.1 Simoa® Kit; Aβ42, Aβ40, and total-tau using the Neurology 3-Plex A Advantage Simoa® Kit, both run on the Simoa® SR-X platform (all Quanterix, USA). All samples were analyzed in duplicate in accordance with the manufacturer’s instructions and standard procedures. All samples from each subject were assayed on the same 96-well-plate within a single run. One participant’s p-tau181 values fell below the quantification limit at two timepoints (S3, S9) and were excluded. Aβ42 and total-tau were log-transformed prior to analysis due to non-normality (Shapiro-Wilk, p < 0.05); log-transformation successfully normalized both distributions. The Aβ42/40 ratio was examined for the six-night comparison only, as prior evidence indicates that Aβ42 and Aβ40 show parallel overnight trajectories whose individual changes largely cancel out when expressed as a ratio (29); analysis of Aβ42/40 ratio also allows direct comparison with Wunderlin et al. (2026).

To control for potential confounding influences of renal and hepatic function on plasma biomarker levels, relevant markers were assessed from a single baseline blood draw (see Supplementary Methods for details).

### Sleep EEG recording and analysis

Full polysomnographic EEG (58 active electrodes; Brain Products actiCAP; 500 Hz) was recorded during S2- S4; a six-electrode reduced montage was used for S5 - S9. Sleep staging was performed offline using the YASA algorithm (30) on 30-second epochs and verified by an expert scorer. Stimulation epochs were not scored due to strong artefacts in the EEG signal, a method also applied to the corresponding sham epochs.

Preprocessing and analysis of EEG data was conducted with MNE-python (v1.7.1) (31), using custom scripts based on the open-source code by Jevri Hanna (https://github.com/jshanna100/sfb/tree/master/sfb2), which we adapted for our needs (https://github.com/jbkordass/memory-sfb2-2025). After applying a 50 Hz notch filter and its harmonics, data was downsampled to 200 Hz, followed by bad-channel identification (ANOAR package) (32) and visual artifact rejection.

EEG analyses focused on 1-minute intervals following each tDCS or sham block at fronto-central derivations (Fz, or FC1/FC2 in case of noise), with a 30-second buffer period. Spectral power analysis was performed using YASA (30) to compute power spectral density (PSD) via Welch’s method on 4-second epochs (50% overlap; Hamming window; frequency resolution 0.25 Hz), with absolute band power calculated for the SO (0.5- 1 Hz) frequency range. SO events were detected following established criteria (27,33), using a 0.16-1.25 Hz finite impulse response bandpass filter; events were retained when peak-to-trough amplitude exceeded the 65th percentile threshold and duration fell between 0.8 and 2 s. Time-frequency representations (TFRs) were computed for each SO-locked epoch using Morlet wavelet convolution (5 cycles; 10- 20 Hz in 0.2-Hz steps) with z-score baseline correction. SO-spindle coupling was assessed via phase-amplitude coupling analyses, with coupling strength (resultant vector length) and mean phase angle (coupling phase) compared across conditions using Watson-Williams tests and repeated-measures ANOVA (Tensorpac 0.6.5).

### Statistical analysis

Analyses were performed in Python (NumPy, pandas, pycircstat2) and SPSS (v29.0). Given the exploratory nature of this pilot study, inference focused on effect size estimation rather than null hypothesis significance testing. Group-level comparisons used paired t-tests and repeated-measures ANOVAs, with Cohen’s *d* (95% CI) and partial η² (95% CI) as effect size metrics, respectively. Stimulation-induced biomarker change was quantified as difference scores (so-tDCS minus sham). Associations between sleep and biomarker changes were examined using linear regression. Although SO-spindle coupling phase angles are inherently circular measures, the observed distribution was tightly confined to a narrow arc of less than 95° (range: −59° to +12° in all sleep sessions), eliminating wrap-around discontinuities; phase angles were therefore treated as linear variables. Model fit was evaluated using R², and adjusted R² is reported as a bias-corrected estimate of explained variance that accounts for sample size and the number of predictors in the model. Where covariates were included (e.g. baseline eGFR), adjusted β coefficients and 95% CIs are reported.

## Results

### Acute effects of a single so-tDCS night on sleep microstructure

Group-level effects of a single so-tDCS night on NREM sleep microstructure are summarized in Table 2. Stimulation was associated with small-to-moderate increases in SO power and SO-spindle coupling strength relative to sham (Cohen’s *d* = 0.33, 95% CI: −0.32 to 0.96, and *d* = 0.47, 95% CI: −0.20 to 1.11, respectively). For SO-spindle coupling phase, the estimated effect size was smaller (η²ₚ = 0.16; 95% CI: 0.00 to 0.43), with coupling timing shifting descriptively toward the SO up-state under so-tDCS (−21.89° ± 3.06°) compared with sham (−31.40° ± 4.05°, Fig. 2).

**Figure 2.**
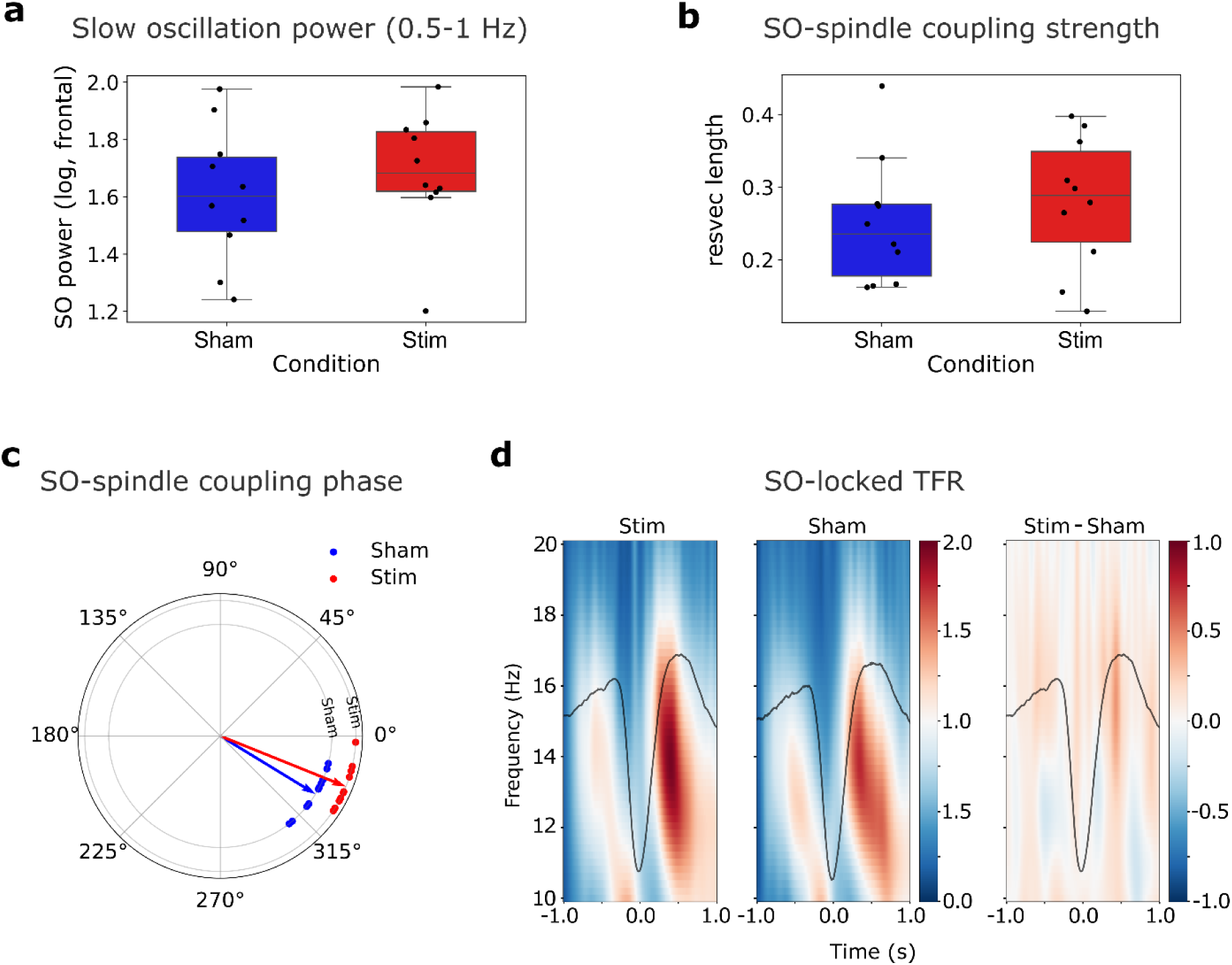
Effects of a single so-tDCS night on NREM sleep microstructure. (a) Slow oscillation (SO) power (0.5-1 Hz, log-transformed, frontal) under sham (blue) and stimulation (red) conditions. (b) SO-spindle coupling strength, quantified as resultant vector length, for sham and stimulation conditions. (c) Polar plot of SO-spindle coupling phase for individual participants (dots) and group mean vectors under sham (blue) and stimulation (red), showing a descriptive shift toward the SO up-state under so-tDCS. (d) SO-locked time-frequency representations (TFR) for the stimulation (left) and sham (middle) conditions, and their difference (Stim − Sham; right), with the average SO waveform overlaid (black line). Warm colors indicate increased power relative to baseline. Boxplots show median, interquartile range, and 1.5× IQR whiskers; individual data points are overlaid.

**Table 2.**
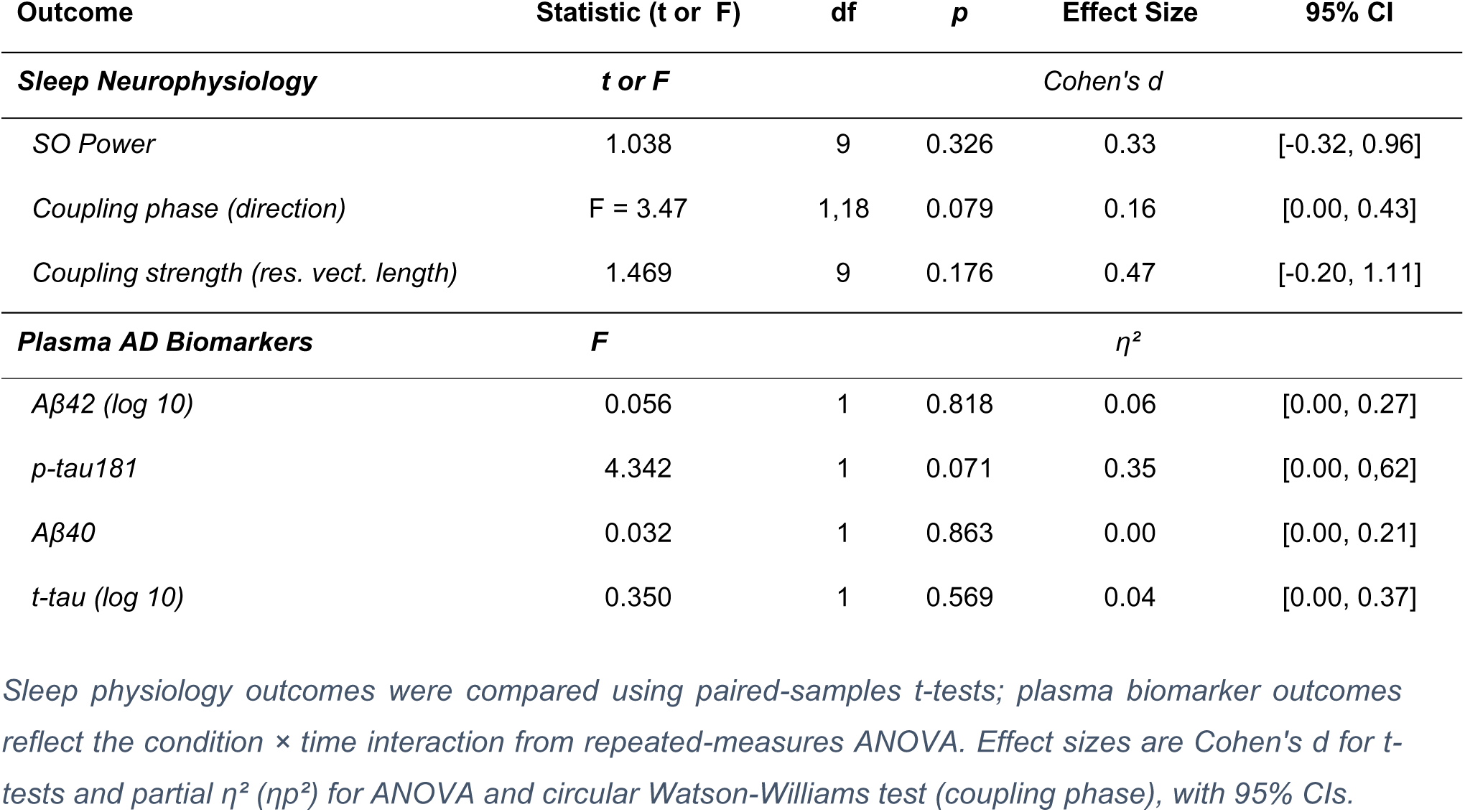
Results of a single so-tDCS night on sleep physiology and AD plasma biomarkers.

In line with these findings, time-frequency analysis showed greater spindle activity (12-16 Hz) during the late rising phase of the SO under stimulation compared to sham (Fig. 2).

### Acute effects on overnight AD plasma biomarker changes

Overnight changes in plasma biomarkers following one so-tDCS night versus one sham night are summarized in Table 2.

p-tau181 showed the largest estimated effect (η²ₚ = 0.35; 95% CI: 0.00- 0.62), with mean overnight plasma levels increasing after so-tDCS (+22.46% ± 13.97%) and slightly decreasing after sham (−7.07% ± 24.17%; Fig. 3).

**Figure 3.**
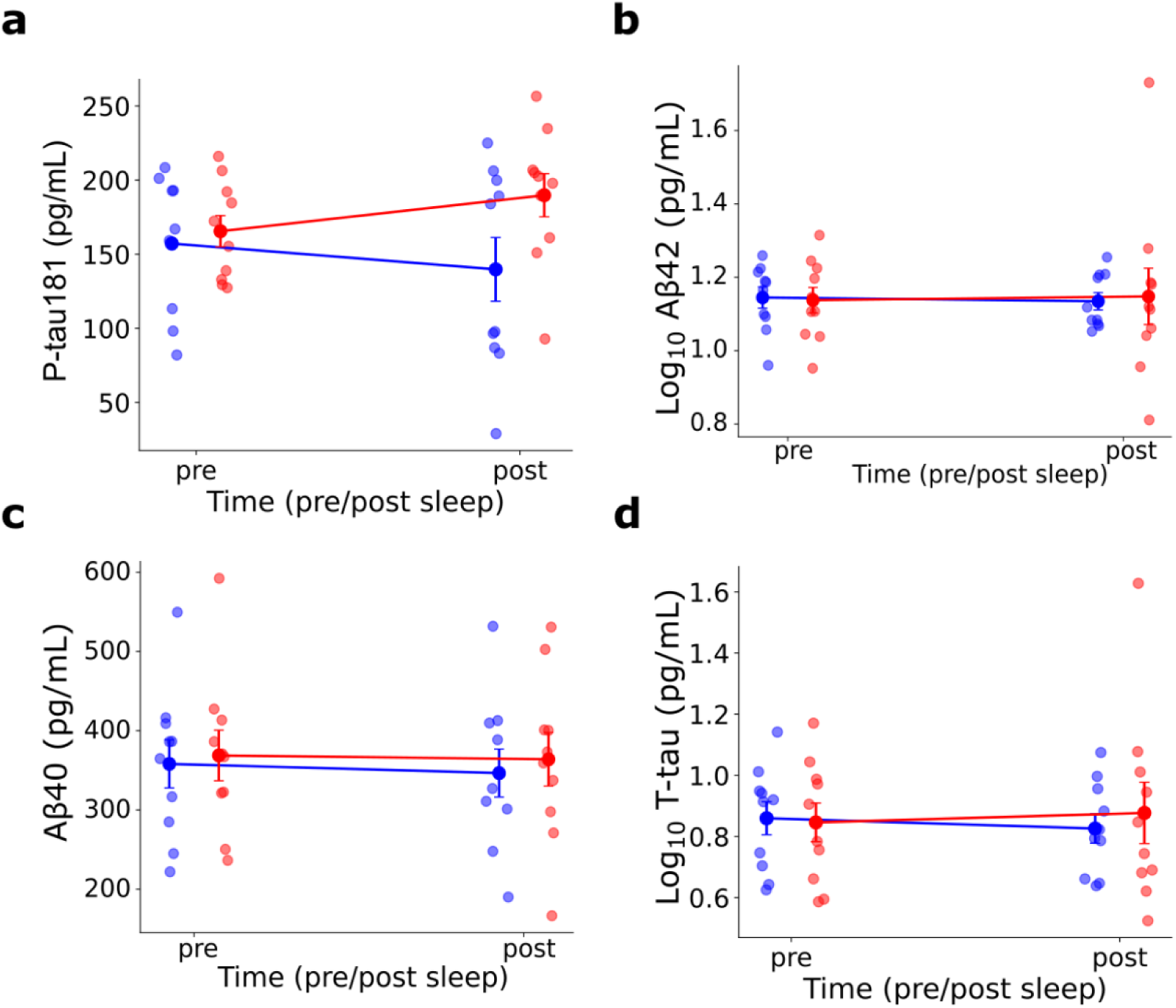
Overnight changes in plasma Alzheimer’s disease biomarkers following so-tDCS versus sham. Pre- and post-sleep plasma levels of (a) phosphorylated tau 181 (p-tau181), (b) β-amyloid 42 (Aβ42, log10-transformed), (c) β-amyloid 40 (Aβ40), and (d) total tau (t-tau, log10-transformed) for the stimulation (red) and sham (blue) conditions. Individual data points are shown as dots; group means ± standard error are depicted as filled circles with error bars. Lines connect group means across timepoints. p-tau181 showed the largest condition difference, with levels increasing overnight following so-tDCS, while Aβ42, Aβ40, and t-tau showed negligible condition differences.

Other biomarkers showed small to at moderate estimated effects with broad confidence intervals (Aβ42: η²ₚ = 0.06, CI up to 0.27; t-tau: η²ₚ = 0.04, CI up to 0.37), while Aβ40 showed a negligible estimated effect (η²ₚ = 0.00; CI up to 0.21).

### Associations between stimulation-related changes in sleep microstructure and overnight biomarker dynamics after one night

To examine whether individual differences in stimulation-induced SO enhancement predicted overnight biomarker changes, difference scores (so-tDCS minus sham) were computed for SO power and each biomarker (i.e., condition differences in the pre- to post-sleep change), and subsequently entered into linear regression models (see Table 3).

**Table 3.** Regressions for acute effects (overnight biomarker changes)

| Predictor | $\beta$ | $R^2$ (95% CI) | Adj. $R^2$ | p | $\beta$<br>(adj. for eGFR) | Adj. $R^2$<br>(adj. for eGFR) | ΔAdj. $R^2$ | p<br>(adj. for eGFR) |
| --- | --- | --- | --- | --- | --- | --- | --- | --- |
| ***p-tau181*** |  |  |  |  |  |  |  |  |
| SO power | 0.76 | 0.58 [0.01, 0.87] | 0.51 | 0.018 | 0.781 | 0.76 | 0.25 | 0.006 |
| Coupling phase | 0.17 | 0.03 [0.00, 0.43] | -0.11 | 0.658 | — | — | — | — |
| Coupling strength | 0.34 | 0.11 [0.00, 0.56] | -0.01 | 0.376 | — | — | — | — |
| ***Log Aβ42*** |  |  |  |  |  |  |  |  |
| SO power | -0.42 | 0.17 [0.00, 0.62] | 0.07 | 0.230 | 0.72 | 0.27 | 0.20 | 0.065 |
| Coupling phase | 0.65 | 0.42 [0.00, 0.78] | 0.35 | 0.043 | — | — | — | — |
| Coupling strength | 0.39 | 0.15 [0.00, 0.60] | 0.05 | 0.261 | — | — | — | — |
| ***Aβ40*** |  |  |  |  |  |  |  |  |
| SO power | -0.47 | 0.22 [0.00, 0.66] | 0.12 | 0.171 | — | — | — | — |
| Coupling phase | 0.63 | 0.39 [0.00, 0.74] | 0.32 | 0.052 | 0.82 | 0.37 | 0.05 | 0.032 |
| Coupling strength | 0.28 | 0.08 [0.00, 0.52] | -0.03 | 0.427 | — | — | — | — |
| ***Log t-tau*** |  |  |  |  |  |  |  |  |
| SO power | -0.41 | 0.17 [0.00, 0.62] | 0.07 | 0.235 | — | — | — | — |
| Coupling phase | 0.53 | 0.28 [0.00, 0.70] | 0.19 | 0.114 | — | — | — | — |
| Coupling strength | 0.37 | 0.14 [0.00, 0.59] | 0.03 | 0.288 | — | — | — | — |

$\beta$ = standardized coefficient. $R^2$ values are reported with 95% confidence intervals (CI). p-values are reported for completeness and interpreted cautiously given the small sample size. $\beta$ (adj. for eGFR) reflects the SO power coefficient after inclusion of renal function as a covariate; Adj. $R^2$ (adj. for eGFR) and p (adj. for eGFR) refer to the full covariate-adjusted model. $\Delta$ Adj. $R^2$ reflects the change in adjusted $R^2$ after inclusion of renal function as a covariate (covariate impact). “—” indicates that no covariate-adjusted model was estimated for that predictor-outcome pair.

Stimulation-induced change in SO power showed a positive association with differences in overnight p-tau181 between conditions, accounting for a substantial proportion of variance (*R*² = 0.58, adj. *R*² = 0.51, 95% CI [0.02, 0.87]; Fig. 4). Greater increases in SO power under so-tDCS relative to sham were associated with larger overnight increases in p-tau181 after so-tDCS relative to sham. In a covariate-adjusted model additionally including baseline kidney function (eGFR), the SO power coefficient remained stable in direction and magnitude (adj. β = 401.37, *t*(8) = 4.48, *R*² = 0.82, adj. *R*² = 0.76, 95% CI [0.19, 0.94]), with SO power accounting for 58.5% and eGFR an additional 24.3% of variance, pointing to partly independent contributions. No meaningful associations were observed between SO power changes and other plasma biomarkers Aβ42 (adj. *R*² ≤ - 0.01-0.12). Additional regression analyses examined associations with SO-spindle coupling and plasma biomarker changes. Stimulation-related changes in coupling strength showed consistently negligible associations with all biomarkers (adj. *R*² ≤ 0.05 - 0.08). In contrast, stimulation-related shifts in SO-spindle coupling phase were substantially associated with both Aβ42 (R² = 0.42, adj. R² = 0.35, 95% CI [0.00, 0.79]) and Aβ40 (R² = 0.39, adj. R² = 0.32, 95% CI [0.00, 0.78]; Fig. 4), but not with p-tau181 or total tau (adj. R² = −0.11 and 0.03, respectively). Specifically, greater alignment of spindle maxima toward the SO up-state under so-tDCS was associated with larger overnight increases in both Aβ isoforms. These associations remained stable after additional eGFR adjustment (Aβ42: adj. R² = 0.27; Aβ40: adj. R² = 0.37).

**Figure 4.**
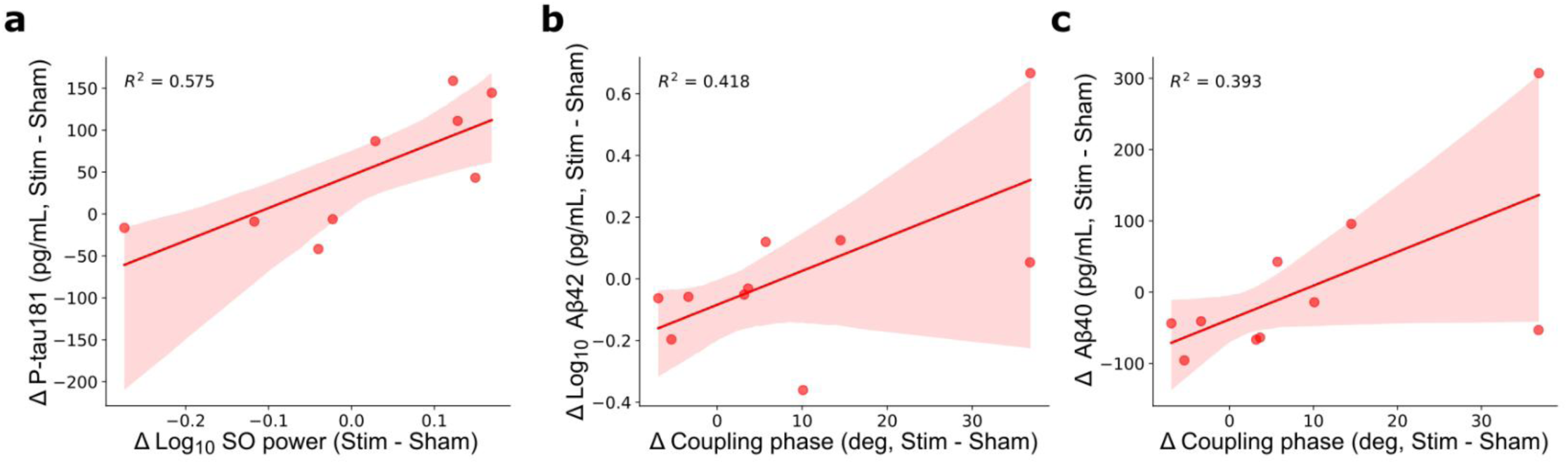
Associations between stimulation-induced changes in sleep microstructure and overnight plasma biomarker dynamics. (a) Stimulation-related change in frontal SO power (log10, Δ stim − sham) positively predicted the condition difference in overnight p-tau181 change, such that greater SO power enhancement under so-tDCS was associated with larger overnight increases in p-tau181. (b–c) Stimulation-related shifts in SO-spindle coupling phase (Δ stim − sham, expressed as mean phase difference) positively predicted condition differences in overnight Aβ42 (log10) and Aβ40, with greater phase alignment of spindle timing toward the SO up-state under so-tDCS associated with larger overnight increases in both amyloid isoforms. Each data point represents one participant. Solid lines show ordinary least-squares regression fits; shaded areas indicate 95% confidence intervals.

Taken together, a single so-tDCS night was associated with small-to-moderate group-level enhancements in sleep microstructure and changes in overnight plasma biomarker levels relative to sham. Notably, participants who showed greater SO power increases under so-tDCS also showed larger overnight increases in p-tau181, an association that explained over half of the between-subject variance and remained stable after adjusting for baseline kidney function. Shifts in SO-spindle coupling phase additionally showed moderate associations with overnight Aβ42 and Aβ40 dynamics, whereas coupling strength and other biomarkers showed negligible associations throughout.

### Multi-night stimulation effects (six nights)

#### Effects on sleep microstructure across six nights

Group-level stimulation effects on NREM sleep microstructure across the six nights are summarized in Table 4. Estimated effects for SO power, coupling strength and coupling phase remained in the small-to-moderate range (d = 0.27- 0.42).

**Table 4.** Results for multi-night so-tDCS effects on sleep physiology and AD plasma biomarkers.

| Outcome | Statistic (t) | df | *p* | Effect Size ($\eta^2$) | 95% CI |
| --- | --- | --- | --- | --- | --- |
| ***Sleep Neurophysiology*** |  |  |  | *Cohen's d* |  |
| *SO Power* | 0.854 | 9 | 0.415 | 0.27 | $[-0.37, 0.90]$ |
| *SO-spindle coupling (phase)* | 0.927 | 9 | 0.378 | 0.29 | $[-0.34, 0.82]$ |
| *Coupling strength (res. vect. length)* | 1.333 | 9 | 0.215 | 0.42 | $[-0.24, 1.1]$ |
| ***Plasma AD Biomarkers (condition × time interaction)*** |  |  |  | *Cohen's d* |  |
| *Aβ42 (log 10)* | -.598 | 9 | 0.564 | -0.19 | $[-0.81, 0.442]$ |

| Outcome | Statistic (t) | df | p | Effect Size ($\eta^2$) | 95% CI |
| --- | --- | --- | --- | --- | --- |
| p-tau181 | 1.411 | 8 | 0.196 | 0.47 | [-0.23, 1.15] |
| Aβ40 | 1.074 | 9 | 0.311 | 0.34 | [-0.31, 0.97] |
| t-tau (log 10) | 0.255 | 9 | 0.805 | 0.08 | [-0.54, 0.67] |

*Sleep physiology outcomes were compared to the sham condition using averaged values across stimulation nights by means of paired-samples t-tests. Cumulative effects on plasma biomarkers were evaluated by comparing morning plasma levels at baseline (T1) and after six stimulation nights (post-intervention, T9) using paired-samples t-tests. Effect sizes are Cohen's d for t-tests with 95% CIs.*

#### Plasma biomarkers

To examine longer-term effects, morning plasma biomarker levels were compared between baseline (S1) and post-intervention (S9, after six stimulation nights). Estimated effects ranged from negligible to moderate (Cohen’s *d* = −0.19 to 0.47; Table 4). Most notably, p-tau181 showed a moderate estimated effect (*d* = 0.47, 95% CI [-0.23, 1.15]), with the point estimate indicating a meaningful tendency toward post-intervention increases. This finding is consistent with the acute overnight p-tau181 response observed after a single so-tDCS night. Aβ40 showed a small-to-moderate estimated effect (*d* = 0.34, 95% CI [−0.31, 0.97]), while t-tau and Aβ42 showed negligible estimated effects (*d* = 0.08, 95% CI [−0.54, 0.67] and *d* = −0.19, 95% CI [−0.81, 0.44], respectively).

**Table 4.** Regressions for longer-term effects.

| Predictor | $\beta$ | $R^2$ (95% CI) | Adj. $R^2$ | p | $\beta$<br>(adj. for<br>eGFR) | Adj. $R^2$<br>(adj. for<br>eGFR) | ΔAdj. $R^2$ | p<br>(adj. for<br>eGFR) |
| --- | --- | --- | --- | --- | --- | --- | --- | --- |
| ***p-tau181*** |  |  |  |  |  |  |  |  |
| SO power | 0.36 | 0.13 [0.00, 0.62] | 0.01 | 0.340 | — | — | — | — |
| Coupling phase | -0.30 | 0.09 [0.00, 0.58] | -0.04 | 0.435 | — | — | — | — |
| Coupling strength | 0.61 | 0.38 [0.00, 0.78] | 0.29 | 0.080 | 0.55 | 0.53 | 0.24 | 0.067 |
| ***Log<sub>10</sub> Aβ42*** |  |  |  |  |  |  |  |  |
| SO power | 0.29 | 0.09 [0.00, 0.53] | -0.03 | 0.411 | — | — | — | — |
| Coupling phase | -0.72 | 0.52 [0.02, 0.84] | 0.46 | 0.018 | -0.83 | 0.56 | 0.04 | 0.009 |
| Coupling strength | 0.32 | 0.10 [0.00, 0.55] | -0.01 | 0.363 | — | — | — | — |
| ***Aβ40*** |  |  |  |  |  |  |  |  |
| SO power | 0.08 | 0.01 [0.00, 0.35] | -0.12 | 0.837 | — | — | — | — |
| Coupling phase | 0.20 | 0.04 [0.00, 0.48] | -0.08 | 0.577 | — | — | — | — |
| Coupling strength | 0.02 | 0.00 [0.00, 0.19] | -0.12 | 0.950 | — | — | — | — |
| ***Log<sub>10</sub> Aβ42/40*** |  |  |  |  |  |  |  |  |
| SO power | 0.24 | 0.06 [0.00, 0.50] | -0.06 | 0.512 | — | — | — | — |
| Coupling phase | -0.74 | 0.55 [0.03, 0.85] | 0.49 | 0.015 | -0.80 | 0.59 | 0.04 | 0.016 |
| Coupling strength | 0.27 | 0.08 [0.00, 0.53] | -0.04 | 0.443 | — | — | — | — |
| ***Log<sub>10</sub> T-Tau*** |  |  |  |  |  |  |  |  |
| SO power | -0.12 | 0.00 [0.00, 0.03] | -0.13 | 0.973 | — | — | — | — |
| Coupling phase | 0.14 | 0.02 [0.00, 0.42] | -0.10 | 0.699 | — | — | — | — |
| Coupling strength | 0.37 | 0.14 [0.00, 0.59] | 0.03 | 0.291 | — | — | — | — |

$\beta$ = standardized coefficient. $R^2$ values are reported with 95% confidence intervals (CI). $p$ -values are reported for completeness and interpreted cautiously given the small sample size. $\beta$ (adj. for eGFR) reflects the SO power coefficient after inclusion of renal function as a covariate; Adj. $R^2$ (adj. for eGFR) and $p$ (adj. for eGFR) refer to the full covariate-adjusted model. $\Delta$ Adj. $R^2$ reflects the change in adjusted $R^2$ after inclusion of renal function as a covariate (covariate impact). “—” indicates that no covariate-adjusted model was estimated for that predictor-outcome pair.

### Associations between multi-night sleep changes and biomarker changes

Average stimulation-related SO power change across multiple nights, compared to sham, showed no meaningful association with baseline-to-post-intervention change in p-tau181 (adj. R² = 0.01, 95% CI [0.00, 0.62], see Table 4). By contrast, average changes in SO-spindle coupling phase relative to sham were substantially associated with Aβ42 change (R² = 0.52, adj. R² = 0.46, 95% CI [0.02, 0.84]; Fig. 5), with a substantial association observed in the covariate-adjusted model including baseline eGFR (R² = 0.66, adj. R² = 0.56, 95% CI [0.01, 0.88]), with eGFR explaining an additional 10% of variance (ΔR² = 0.10). A similar pattern was observed for Aβ42/40 ratio (R² = 0.55; covariate-adjusted model R² = 0.59), while no associations were observed for Aβ40 and t-tau with SO-spindle coupling phase (see Table 4 for more details).

**Figure 5.**
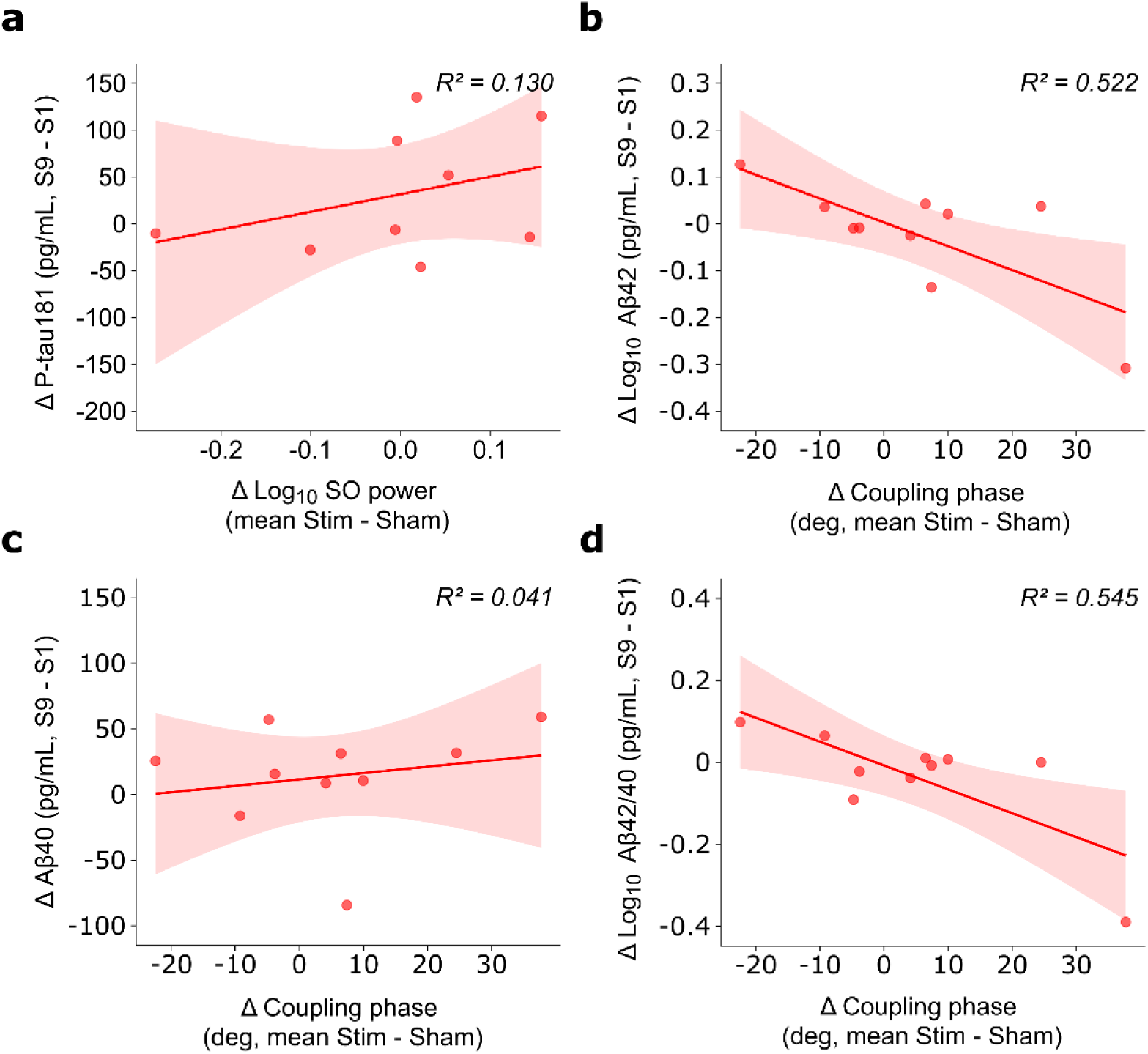
Associations between multi-night stimulation-induced changes in sleep microstructure and long-term plasma biomarker changes. (a) Average stimulation-related change in SO power across multiple nights showed no meaningful association with baseline-to-post-intervention change in p-tau181. (b) Mean stimulation-related shift in SO-spindle coupling phase (average of T3 to T9, Δ stim − sham) showed a negative association with change in Aβ42 (log10), whereby greater phase alignment toward the SO up-state was associated with greater Aβ42 reduction over the intervention period. (c) No meaningful association was observed between mean coupling phase shift and Aβ40 change. (d) Coupling phase shift showed a negative association with change in the Aβ42/40 ratio (log10), similar to Aβ42 (log10). Each data point represents one participant. Solid lines show ordinary least-squares regression fits; shaded areas indicate 95% confidence intervals.

### Consistency of individual plasma biomarker responses to so-tDCS across time scales

Finally, we evaluated whether between-person variability in the acute overnight plasma change aligned with change across the six-night intervention: we examined the association between overnight plasma biomarker changes following a single stimulation night and the baseline-to-post-intervention change (morning levels). Overnight p-tau181 change showed a substantial positive association with baseline-to-post-intervention change (R² = 0.69, adj. R² = 0.64), indicating that participants who exhibited larger acute overnight increases also tended to show larger increases over the intervention period. For Aβ42, a moderate negative association was observed (R² = 0.28, adj. R² = 0.19, 95% CI [0.00, 0.72]), such that participants with larger acute overnight Aβ42 increases tended to show smaller baseline-to-post-intervention increases, a pattern opposite to that seen for p-tau181 and for the association between overnight SO-spindle coupling phase and Aβ42. No such consistency was found for Aβ40 (adj. R² = −0.01). Notably, overnight change in Aβ42 showed a substantial negative association with baseline-to-post-intervention change in the Aβ42/40 ratio (R² = 0.63, adj. R² = 0.58), such that participants with greater overnight Aβ42 increases after stimulation tended to show larger decreases in the Aβ42/40 ratio across the intervention period.

Taken together, six consecutive so-tDCS nights produced small-to-moderate enhancements in sleep microstructure on average, while group-level plasma biomarker effects remained negligible to small. A notable exception was p-tau181, which showed a moderate tendency toward post-intervention increases, consistent with the acute findings. Unlike the single-night results, however, mean individual SO power changes across the six nights did not predict longer-term p-tau181 trajectories. Instead, SO-spindle coupling phase shifts toward the SO up-state emerged as the stronger longitudinal predictor, associating with decreases in both Aβ42 and the Aβ42/40 ratio.

## Discussion

This exploratory mechanistic study in healthy older adults provides preliminary evidence that modulation of sleep slow oscillatory dynamics via so-tDCS is associated with changes in peripheral AD-related biomarkers. Distinct patterns of association emerged for different features of sleep microstructure, with partially dissociable relationships observed for SO power and SO-spindle coupling. While group-level stimulation effects on sleep dynamics and plasma biomarkers were small to moderate, at the inter-individual level, stimulation-induced increases in SO power showed a strong positive association with overnight changes in p-tau181. SO-spindle coupling phase was moderately associated with overnight increases in Aβ42 and Aβ40, as well as longer-term decreases in Aβ42 and Aβ42/40 ratio across the multi-night intervention.

### Slow oscillation activity and p-tau181: signals of inter-compartmental redistribution

The most robust finding of this study was a strong positive association between stimulation-induced increases in SO power and overnight increases in plasma p-tau181 relative to sham, which accounted for over half of the between-subject variance and remained stable after statistical adjustment for baseline kidney function (eGFR). This association is consistent with the view that high-amplitude SWA facilitates the mobilization of AD-related proteins from brain interstitial system to CSF and ultimately to peripheral blood via glymphatic-mediated efflux, a process thought to be predominantly active during deeper NREM sleep phases (12,17). Mechanistically, Fultz and colleagues (2019) demonstrated in healthy adults that slow fluctuations in neural activity during NREM sleep were coupled with complementary oscillations in CSF flow as measured by fMRI, providing direct in vivo support for the idea that coordinated cortical slow oscillations drive CSF-interstitial fluid exchange along perivascular pathways. The present data extend prior human evidence linking SWA to biomarker dynamics: whereas earlier studies established that sleep deprivation elevates CSF Aβ and tau levels (3,15,16) and that experimental disruption of SWA acutely elevated overnight CSF Aβ42 concentrations (4), few studies have used an intervention enhancing SWA to examine the resulting biomarker response (20). The direction of the observed p-tau181 increase - rising plasma levels following so-tDCS compared with sham - is consistent with a redistribution rather than a pathological production interpretation: augmented SWA promotes efflux of p-tau from brain parenchyma into CSF and subsequently into peripheral blood, transiently elevating plasma concentrations before peripheral elimination. This aligns with Dagum et al. (2026), who directly tested whether glymphatic clearance during sleep increases morning plasma AD biomarker levels in a multi-site randomized crossover study (sleep vs. sleep deprivation). Using a novel device measuring brain parenchymal resistance to glymphatic flow (Rₚ) by transcranial multifrequency impedance spectroscopy combined with sleep EEG, they found that changes in Rₚ and SWA power predicted overnight changes in plasma Aβ42, p-tau181, as well as np-tau181 and np-tau217. Their multicompartment model further reproduced the preferential clearance of aggregation-prone species such as Aβ42 and p-tau181 relative to their non-aggregation-prone counterparts, consistent with a glymphatic efflux mechanism.

Our experimental manipulation adds causal leverage to this framework: actively driving SO power increases via so-tDCS appears sufficient to induce plasma p-tau181 changes consistent with enhanced glymphatic efflux. The specificity of this association to p-tau181, but not Aβ isoforms and t-tau, may reflect methodological differences to Dagum et al., who used a direct physiological measure of glymphatic resistance integrated over the full night rather than stimulation-induced increments relative to sham. Alternatively, this selectivity may reflect genuine biological specificity, at least within the context of a single stimulation night in cognitively healthy older adults.

### SO-spindle coupling phase and Aβ isoform dynamics

For Aβ dynamics, the temporal precision of SO-spindle coupling, specifically the phase angle at which spindle activity occurred relative to the SO, was the relevant predictive variable. Greater alignment of spindle maxima toward the SO up-state, under so-tDCS relative to sham, was associated with larger overnight increases in Aβ42 and Aβ40 after a single stimulation night (pre-to post sleep change). In contrast, shifts in coupling phase angle across six nights predicted longitudinal decreases in Aβ42 and the Aβ42/40 ratio.

These findings reinforce the idea that the temporal coordination of sleep oscillations carries biologically meaningful information that is not captured by spectral power alone. Tightly coupled SO-spindle interactions reflect the functional integrity of thalamocortical circuits, and their relevance for memory consolidation during sleep is well established (7,27,35). More recently, attention has shifted toward their potential role in the clearance of AD-related proteins. Wunderlin and colleagues found in a sample of healthy and cognitively impaired older adults that SO-spindle coupling consistency was a stronger predictor of baseline plasma Aβ42/40 ratio than SWA power, age, or cognitive functioning (19). Another study by the same group reported that improvements in SO-spindle coupling via acoustic stimulation during sleep were associated with delayed changes in plasma Aβ42/40 ratio in cognitively impaired individuals, but not cognitively healthy older adults (20).

The reason why SO-spindle coupling precision specifically predicted Aβ dynamics remains unclear, and the present data cannot resolve the underlying mechanism. One possibility is that clearance of Aβ preferentially involves passage via meningeal lymphatic vessels to extracranial lymph nodes. This interpretation receives independent support from Eide et al. (2023), who employed two in vivo markers of clearance function, i. e., one proxy of glymphatic clearance, and one proxy of meningeal lymphatic function, to determine their contribution to clearance in sleep deprivation versus natural sleep. They found that overnight changes in plasma Aβ40 and Aβ42 correlated with meningeal lymphatic clearance capacity during sleep for both conditions, while p-tau181 changes correlated with the glymphatic clearance marker in the sleep-deprived group only. Whether SO-spindle coupling precision might influence meningeal lymphatic clearance is unknown, but an indirect pathway is conceivable: spindle activity has been associated with transient cardiac deceleration consistent with increased parasympathetic tone (37), and SO activity with coordinated cerebrovascular oscillations coupled to CSF flow dynamics (17); their precisely timed co-occurrence during sleep may thereby create simultaneously favorable autonomic and hemodynamic conditions for CSF drainage into meningeal lymphatic vessels. This remains speculative and warrants direct experimental investigation.

The mechanism underlying the longitudinal plasma Aβ42 decrease and its apparent divergence from the acute overnight direction remains to be fully elucidated. Notably, the direction of the longitudinal Aβ42/40 change observed here differs from the increase reported by Zeller et al. (20) following acoustic stimulation during three consecutive nights in cognitively impaired older adults, possibly reflecting differences in baseline amyloid burden: in individuals with pathological accumulation, enhanced clearance is thought to release plaque-sequestered Aβ42 into the bloodstream, raising the plasma Aβ42/40 ratio (20), whereas in cognitively healthy older adults, any plaque-derived Aβ42 reservoir is expected to be substantially smaller. Thus, participants showing greater stimulation-induced SO-spindle coupling phase shifts, who also showed the largest longitudinal Aβ42 decreases, may have benefited from more consistently enhanced nightly clearance, progressively reducing the cerebral ISF and CSF Aβ42 pool toward a lower steady-state and thereby reducing the net flux of Aβ42 into the systemic circulation over successive nights. This hypothesis remains to be assessed with parallel measurements of cerebral and peripheral Aβ42 dynamics.

### Implications for glymphatic and multi-component clearance models

The present findings extend the observational framework established by Dagum et al. (2026), who showed that naturally occurring variation in SWA (power) and glymphatic resistance predicts plasma biomarker changes during sleep. By demonstrating that experimentally induced increments in SO power are linked to p-tau181 dynamics in a dose-response-like fashion at the individual level, the present study lends support to the inference that SWA plays a mechanistic rather than merely correlational role in p-tau clearance, and that non-invasive brain stimulation can modulate the mobilization of AD-relevant proteins from brain to blood.

The dissociation between SO power and SO-spindle coupling phase effects on p-tau181 versus Aβ dynamics is furthermore broadly consistent with a multi-compartment redistribution framework in which distinct features of sleep physiology contribute to different steps of the overall clearance pathway (34,36). It further implies that comprehensively targeting both p-tau and Aβ may require to optimize both SO amplitude and the temporal precision of SO-spindle coupling.

Finally, the present findings extend prior work with acoustic closed-loop stimulation (20) to transcranial electrical stimulation, and demonstrate sleep-dependent biomarker modulation in cognitively healthy older adults rather than exclusively in those with cognitive impairment.

## Limitations

Some limitations of the study should be noted. First, the sample size was modest, which limits the precision of effect estimates. Confidence intervals are correspondingly wide, and results should be treated as exploratory rather than confirmatory. Second, we relied on plasma biomarkers as indirect proxies of central nervous system clearance rather than directly measuring Aβ or tau dynamics in CSF or via PET imaging. Plasma measures provide a scalable, sensitive, and minimally invasive index of AD-related biology, though plasma levels are influenced by peripheral production, metabolism, and clearance pathways (38). To mitigate this, we implemented standardized fasting and sampling procedures and statistically adjusted for renal function. Third, in our sample of cognitively healthy older adults, CSF or PET biomarkers were not available to exclude AD pathology.

## Conclusion

This study provides initial evidence that non-invasive enhancement of slow oscillatory dynamics during sleep modulates peripheral AD biomarkers in healthy older adults, with dissociable links between SO power and p-tau181 dynamics on the one hand, and SO-spindle coupling precision and Aβ dynamics on the other. Given that sleep disturbances precede clinical AD by decades and that SWA declines progressively with normal aging, these findings are relevant to the development of early preventive strategies targeting the sleep-dependent clearance of AD-related proteins before pathology becomes irreversible. They further suggest that optimizing both SO amplitude and SO-spindle coupling, rather than SWA alone, may be necessary to comprehensively address tau and Aβ dynamics, and provide effect-size estimates to inform future confirmatory trial designs.

## Availability of data and materials

The data underlying this article will be shared on reasonable request to the corresponding author.

## Acknowledgments

We thank M. Johns for her vital contribution, and all participants for taking part in the study.

## Author contributions

J.L., Conceptualization, Design of the work, Funding acquisition, Interpretation of Data, Supervision, Project administration, Formal analysis, Draft, Visualization. P.S., Acquisition, Data curation, Formal analysis, Project administration, Review. Y.R. Acquisition, Data curation, Formal analysis, Project administration, Review. R.M., Formal analysis, Software, Review; B.D. Data curation, Formal analysis, Project administration, Review. A.V. Formal analysis, Review. A.F., Resources, Supervision, Review.

