## Supplementary Methods for "Enhancing slow-wave sleep via non-invasive brain stimulation modulates brain-to-blood clearance of Alzheimer’s disease biomarkers"

### Participants: full exclusion criteria

Participants were screened for the following exclusion criteria: severe untreated medical, neurological, or psychiatric conditions (including neurodegenerative diseases, stroke, epilepsy, depression, and psychosis); intake of medication passing the blood-brain barrier (including antipsychotics, antidepressants, benzodiazepines, and sleep-inducing medications); alcohol or substance abuse; manifest sleep disorders (including sleep apnoea and restless legs syndrome); non-fluency in German; and brain pathologies detected on MRI (including brain tumour and previous stroke).

### Hepatic and renal function assessment

To control for individual differences in peripheral clearance of plasma AD biomarkers, hepatic and renal function markers were assessed from a single fasting blood draw at baseline (S1, 08:00). A total of 3.5 ml of blood was collected in lithium heparin tubes and analyzed at the clinical laboratory of the University Medicine Greifswald. The following markers were assessed: alanine aminotransferase (ALT), aspartate aminotransferase (AST), De Ritis ratio (AST/ALT), creatinine, Cystatin C, and estimated glomerular filtration rate (eGFR), calculated using the CAPA equation (Grubb et al. 2014) and standard estimation methods. The following markers were assessed: alanine aminotransferase (ALT), aspartate aminotransferase (AST), De Ritis ratio (AST/ALT), creatinine, Cystatin C, and estimated glomerular filtration rate (eGFR), calculated using the CAPA equation (1).

For associations reaching at least a moderate effect size, primary models were re-estimated with eGFR included as a covariate in a pre-specified sensitivity analysis to evaluate the robustness of findings against potential confounding by peripheral clearance, consistent with evidence that renal filtration is a relevant determinant of plasma AD biomarker concentrations (2,3).

**EEG Analyses:**

**Slow oscillation (SO) event detection.** The EEG signal was bandpass filtered using a finite impulse response (FIR) filter (0.16-1.25 Hz), and all negative-to-positive zero-crossings were identified. The interval between successive negative zero-crossings was measured, and the peak-to-trough amplitude was computed as the difference between the down-state trough (most negative point following the first positive-to-negative crossing) and the up-state peak (most positive point following the subsequent negative-to-positive crossing). Events were retained if the amplitude difference exceeded the 65th percentile across all detected events in each participant and if the duration between consecutive positive-to-negative zero-crossings was between 0.8 and 2 s, consistent with previously established criteria (4,5).

**Time-frequency representation analysis.** TFRs were computed on 5-second SO-locked epochs (downsampled to 50 Hz) using Morlet wavelet convolution (5 cycles) across 10-20 Hz in steps of 0.2 Hz. Edge effects were minimised by excluding the first and last 150 ms of each epoch. Baseline correction was performed using z-score normalisation relative to the -2.35 to -1.5 s pre-event window.

**SO-spindle coupling.** Phase-amplitude coupling was quantified from SO-locked event segments. The instantaneous SO phase angle was extracted by filtering in the 0.5-1.25 Hz band and applying a wavelet transform; spindle amplitude was obtained analogously by filtering in the spindle band (12-15 Hz) followed by wavelet transformation. Analyses were restricted to the -2 to 2 s window to minimize filter edge artefacts. For each participant and SO event, the maximal spindle amplitude and its corresponding SO phase angle were extracted, yielding per-participant distributions of coupling phase angles (with 0° representing the positive peak/up-state and 180° the negative trough/down-state). Mean coupling phase angle and coupling strength (resultant vector length) were derived from these distributions and compared across conditions using Watson-Williams tests (circular statistics) and repeated-measures ANOVA; planned contrasts (cathodal vs. sham; anodal vs. sham) were applied where a significant main effect was observed. All coupling analyses were performed using Tensorpac 0.6.5 (6).
